# Correlated selection on amino acid deletion and replacement in mammalian protein sequences

**DOI:** 10.1101/215277

**Authors:** Yichen Zheng, Dan Graur, Ricardo B. R. Azevedo

## Abstract

A low ratio of nonsynonymous and synonymous substitution rates (dN/dS) at a codon is a sign of functional constraint caused by purifying selection. Intuitively, the functional constraint would also be expected to prevent such a codon from being deleted. Oddly, to the best of our knowledge, the correlation between the rates of deletion and substitution has never actually been estimated. Here, we use 8,595 protein coding-region sequences from 9 mammalian species to examine the relationship between deletion rate and dN/dS. We found significant positive correlations at both the level of sites and genes. We compared our data against controls consisting of simulated coding sequences evolving along identical phylogenetic trees, where the correlation is not included in the model *a priori*. A much weaker correlation was found in the corresponding simulated sequences, which is probably caused by alignment errors. In the real data, the correlations cannot be explained by alignment errors. Separate investigations on nonsynonymous (dN) and synonymous (dS) substitution rates indicate that the correlation is most likely due to a similarity in patterns of selection rather than mutation rates.

## Introduction

The functional constraint on a genomic region is defined by its sensitivity to mutations, that is, the proportion of mutations that negatively affect its function (Graur 2016, pp. 116–120). Genomic regions subject to strong functional constraints are expected to perform important functions and to evolve relatively slowly. Mutations can take many forms, including nucleotide substitutions, insertions, and deletions (indels); as a result, functional constraint can be defined separately with respect to each type of mutation. One might expect functional constraints with respect to nucleotide substitutions and indels to be correlated — if the function of a genomic region can be disrupted by a substitution it can probably also be disrupted by an indel. However, this is not necessarily the case. For example, nucleotide substitutions at a fourfold degenerate site in a protein-coding gene may be selectively neutral because the protein product is not affected. If that fourfold degenerate site is deleted, however, it will cause a frameshift that will likely disrupt the function of that protein severely. Sites that only experience selection when they are deleted were referred to as “indifferent DNA” (Graur et al. 2015).

Substitutions have been studied more extensively than deletions for two reasons. First, because indels are more difficult to detect than substitutions (Landan and Graur 2009; Nagy et al. 2012). Second, because indels have not been modeled mathematically as well as substitutions (but see Lunter et al. 2006). Nevertheless, a few studies have attempted to compare the patterns of functional constraint arising from both kinds of mutations. Taylor et al. (2004) identified 1,743 indel events in 1,282 genes (out of a dataset of 8,148 genes) from human-mouse-rat triple alignments. They compared indel rates in genes of different functions using Gene Ontology (Ashburner et al. 2000), and found that intracellular proteins and enzymes are less likely to have indels. When the indel rate differences were compared with substitution rates (Waterston et al. 2002), a highly similar distribution among categories was found. These results indicate that functional categories that are more “important” to an organism tend to have both reduced amino acid replacement and reduced amino acid loss. One limitation of this study is that it focused on groups of genes rather than on individual genes.

Another study (Miller et al. 2007) used a 28-vertebrate alignment to study coding-sequence conservation. The authors tested the hypothesis that more conserved amino acids are more likely to cause diseases when deleted. They analyzed the gene encoding the enzyme phenylalanine hydroxylase, a gene whose mutations may cause phenylketonuria. The conservation levels of codons involved in disease-causing deletions turned out to be the same as for the gene overall. Miller et al. (2007) concluded that long-term selection against nonsynonymous mutations is consistent with short-term selection (as implied by diseases) against amino acid deletions. One strength of this study was the ability to identify deleterious mutations directly from clinical data. It is, however, only based on a single gene.

Chen et al. (2009) studied the ratio of nucleotide substitution to indel rates, across mammalian and bacterial genomes. They interpreted the ratio as an indicator of the relative strengths of selection on the two types of mutations. They found that, within coding regions, more conserved genes have higher substitution to indel ratios than less conserved genes. This result suggests that indels (even non-frameshifting ones) are subject to relatively stronger selection than substitutions in conserved genes. However, as the comparison is focused on which type of mutation is more common, it does not directly help to resolve the correlation *between* these two types of changes.

In a population-level comparison between 179 human genomes, Montgomery et al. (2013) found that indel-based variations were highly localized: half of them were identified in only ~4% of the genome, likely due to mutation rate effects. The mutation rate heterogeneity was different between indels and substitutions; for example, recombination hotspots accompanied an increase of indels but not SNPs. As expected, the authors found evidence that indels in protein-coding sequences are subject to strong purifying selection. Indeed, even non-frameshift indel variants were found to have lower allele frequencies (a hallmark of purifying selection) than non-coding indels.

From 14 species along the entire tree of life, the evolution of indel rates were analyzed by comparing protein sequences (Sung et al. 2016). It was discovered that indel rate correlates negatively with effective population size, which is already well-known for substitution rates (Lynch 2010). This is consistent with the Drift-Barrier Hypothesis, stating in this case that natural selection is expected to reduce mutation rate to the point where further reduction does not provide enough of a selective benefit to be more likely to fix when compared to a neutral mutation (Sung et al. 2012). However, the selection discussed in that paper is selection *on mutation rate*; it does not directly address the selection against substitution and indels in the entire genome or exome.

While these aforementioned studies addressed the relationship between purifying selection against nonsynonymous substitutions and purifying selection against deletions, they did not fundamentally answer the question: are the two selection effects correlated across the genome, and what is the extent of this correlation? If the correlation does not exist, one can expect the dN/dS ratio of a codon is independent from its deletion rate. If there is a correlation, the dN/dS ratio would be proportional to the probability of the codon being deleted. dN will also behave similarly because it is also under selection, while dS would not because it measures neutral substitutions (Nei and Gojobori 1986; Price and Graur 2016). It is also possible that the correlation occurs only at the gene level, i.e., genes with higher dN/dS would have higher deletion rates, but within a gene, the dN/dS and deletion rate of sites are independent. However, selection is not the only evolutionary force that may cause a correlation between substitutions and indels; it is possible that regions with high point mutation rates would also have high indel mutation rates. In this case, we may see that the deletion rate would be correlated to both dN and dS, but not with the dN/dS ratio.

We used mammalian protein-coding sequences and simulated sequences to study the correlation between deletion rates and dN/dS, to understand how similar the patterns of the two types of selection are. In addition, we used dN and dS separately to estimate their correlations with deletion rates, to test our hypothesis on whether or not mutation plays a role. We have found that there is indeed a positive correlation between the rates of deletion and substitution, and it is likely to be caused by selection, rather than mutation.

## Results

### The deletion and nonsynonymous substitution rates per site are positively correlated

We collected sequences of protein-coding genes from 9 mammalian genomes (Fig. 1, Lindblad-Toh et al. 2011), and aligned them with PROBCONS (Do et al. 2005). Simulated alignments were produced along the same phylogenetic tree with realistic parameters derived from the real data (See Supplementary Text and Fig. S1). In-frame deletions of length 1-8 amino acids were identified (Fig. 2; see Material and Methods for details); deletion rate, dN, dS and dN/dS were measured on each codon. Hereafter, we refer to dN, dS, and dN/dS collectively as “substitution measures.”

**Fig. 1.**
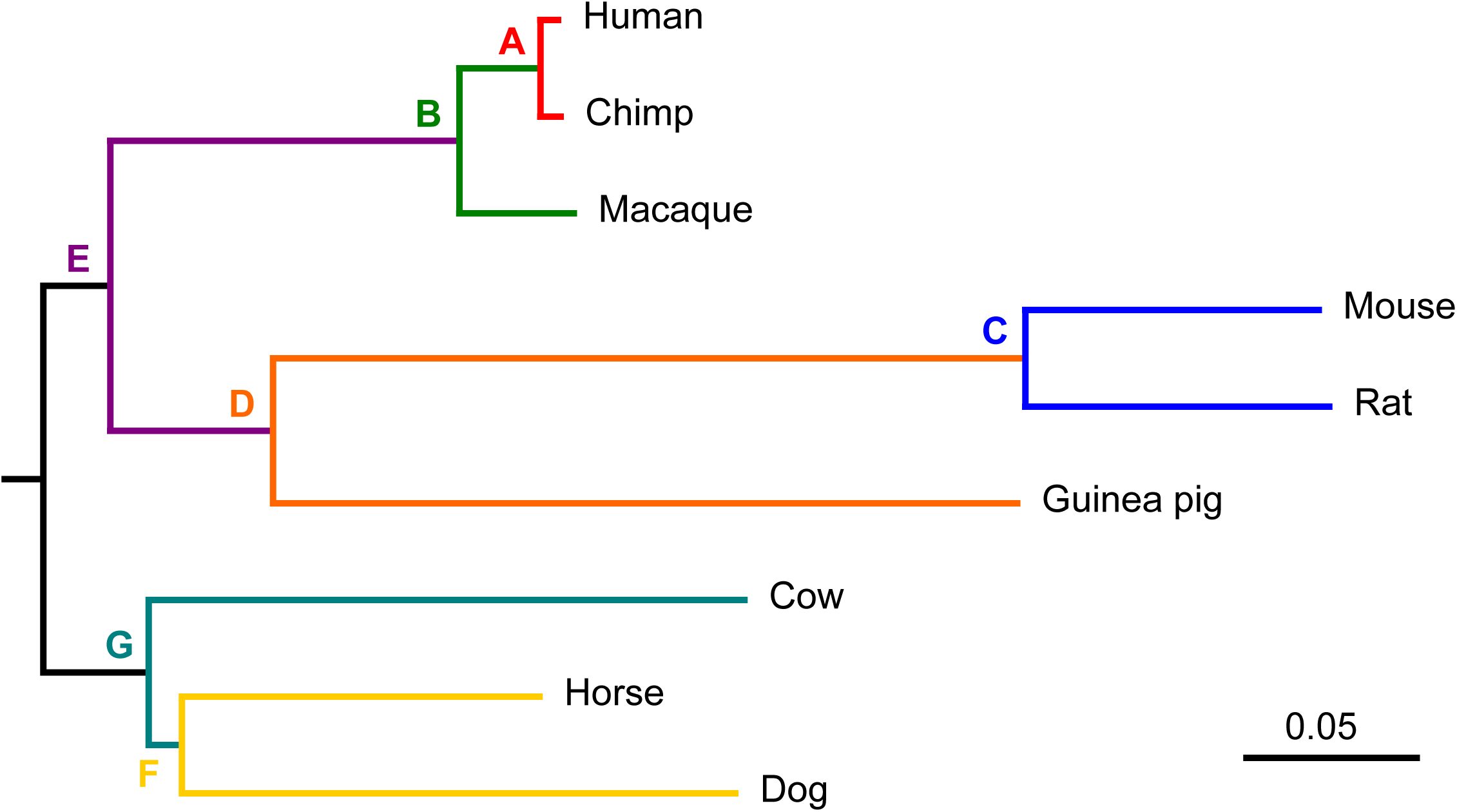
The commonly accepted phylogenetic relationship among the 9 species used in this study. This tree will be called the external reference tree throughout the paper. Seven different colors denote seven pairs of branches/lineages (A–G) on which deletions were estimated. The black-colored branches are the root of the tree. The branch lengths of the 9-species tree are derived from UCSC Human/hg19/GRCh37 46-way multiple alignment (Kent et al. 2002). These branch lengths are used as guidance for simulation and estimation of deletion rates.

**Fig. 2.**
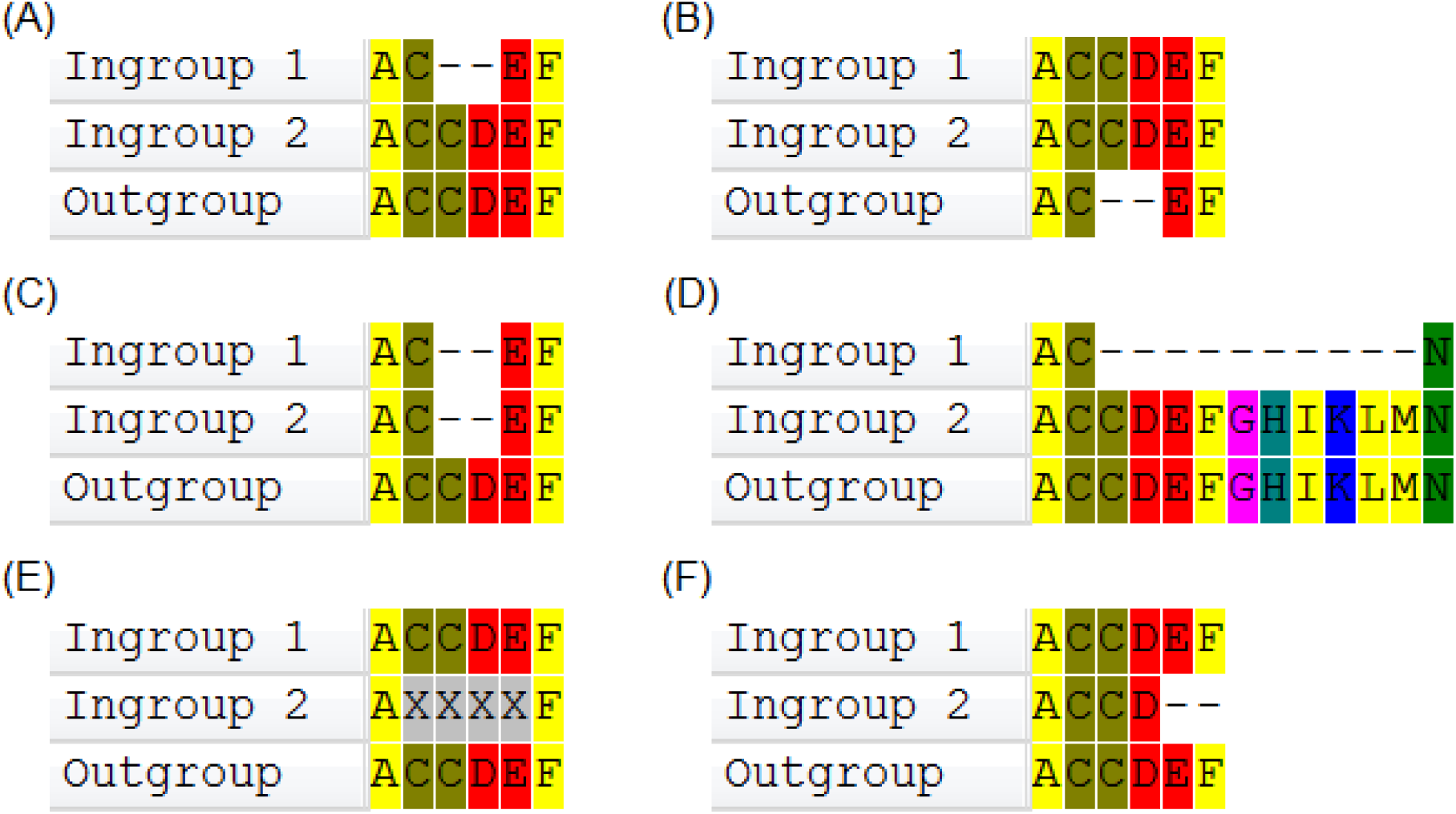
Illustration of how we identify deletion events and non-used sites in protein sequences for each pair of lineages. **A.** Identified short deletion at sites 3 and 4 in taxon 1. **B.** Excluded sites 3 and 4 because of gaps in the outgroup. **C.** Excluded sites 3 and 4 because both ingroup taxa contain gaps at those positions, thus it is impossible to know whether it is an insertion or a deletion. **D.** Excluded sites 3–12 because of long (> 8aa) deletion. **E.** Excluded sites 2–5 because of unknown amino acids. **F.** Excluded sites 5 and 6 because they are included in a terminal gap.

The correlations between deletion rate and the different substitution measures are summarized in Fig. 3. In the “All” dataset the deletion rate is positively correlated with both dN (ρ = 0.11) and the dN/dS ratio (ρ = 0.08) (Fig. 3A). The corresponding correlations in the simulated data are much lower (mean ρ = 0.01 and 0.03 for dN/dS and dN, respectively, based on 1,000 bootstrap replicates; *Z*-test: *Z* = 57.87 for dN/dS and *Z* = 61.02 for dN, *P* < 0.0001 in both cases). The signal is even stronger when the true alignments from simulated data are used, indicating the alignment error causes a small inflation of dN/dS and dN estimates (Fig. S2). The deletion rate is also positively correlated with dS but the correlation is weaker than for dN (ρ = 0.04, Fig. 3A); however, this correlation is significantly stronger in the real data when compared to the simulated data (mean ρ = 0.01; *Z* = 43.36, *P* < 0.0001).

**Fig. 3.**
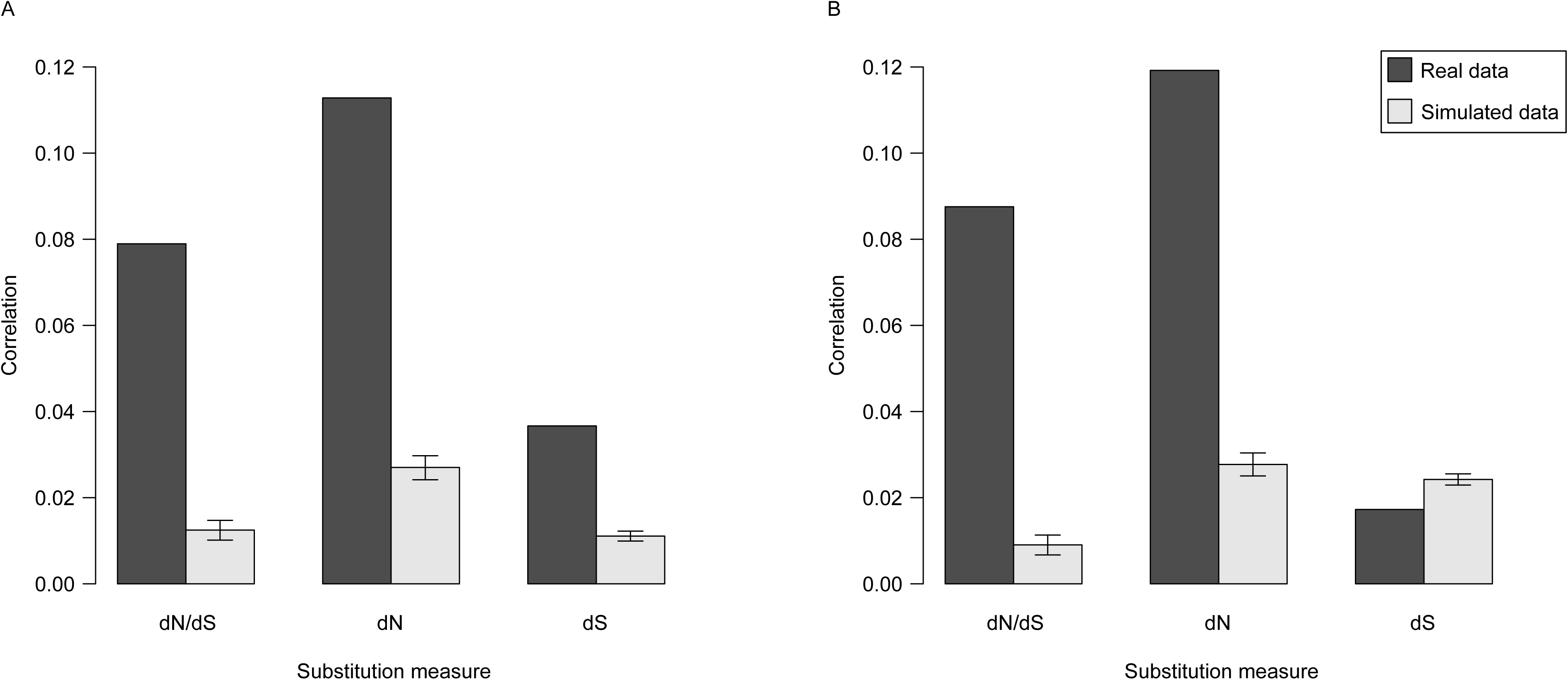
Rates of deletion and substitution per site are positively correlated. Spearman correlation between deletion rate and substitution measures (dN/dS, dN and dS) in real and simulated data. **A.** Based on the “All” dataset. **B.** Based on the “NC-4+” dataset, where all sites without any substitutions or present in less than four species were removed. For the simulated data, the value shown is the mean of 1,000 bootstrap replicates, and the error bars are 2.5% to 97.5% quantiles. Real data produces higher correlations than simulated data for all measures.

To evaluate the robustness of the patterns summarized in Fig. 3A to uncertainty in the estimates of deletion and substitution rates, we repeated the analysis on the “NC-4+” dataset, containing only sites that have at least one nucleotide substitution and that are present (i.e., not gaps) in at least 4 species. The patterns for dN and dN/dS are essentially unchanged (Fig. 3B). However, the correlation between deletion rate and dS disappears; indeed, the correlation is *higher* in the simulated data than in the real data (*Z* = –10.69, *P* < 0.0001). Results for datasets with different thresholds of non-gap characters are similar to those for “NC-4+”. These results indicate that the correlation between deletion rate and dS is largely driven by sites with low substitution rate and/or uncertain estimates of rates of deletion and substitution. We conclude that rates of deletion and nonsynonymous substitution per site are positively correlated. These results indicate that functional constraints against amino acid replacement and against amino acid loss are correlated to each other. Fig. 4 shows the correlation with a density heatmap, where combinations of substitution measures and deletion rate are plotted.

**Fig. 4.**
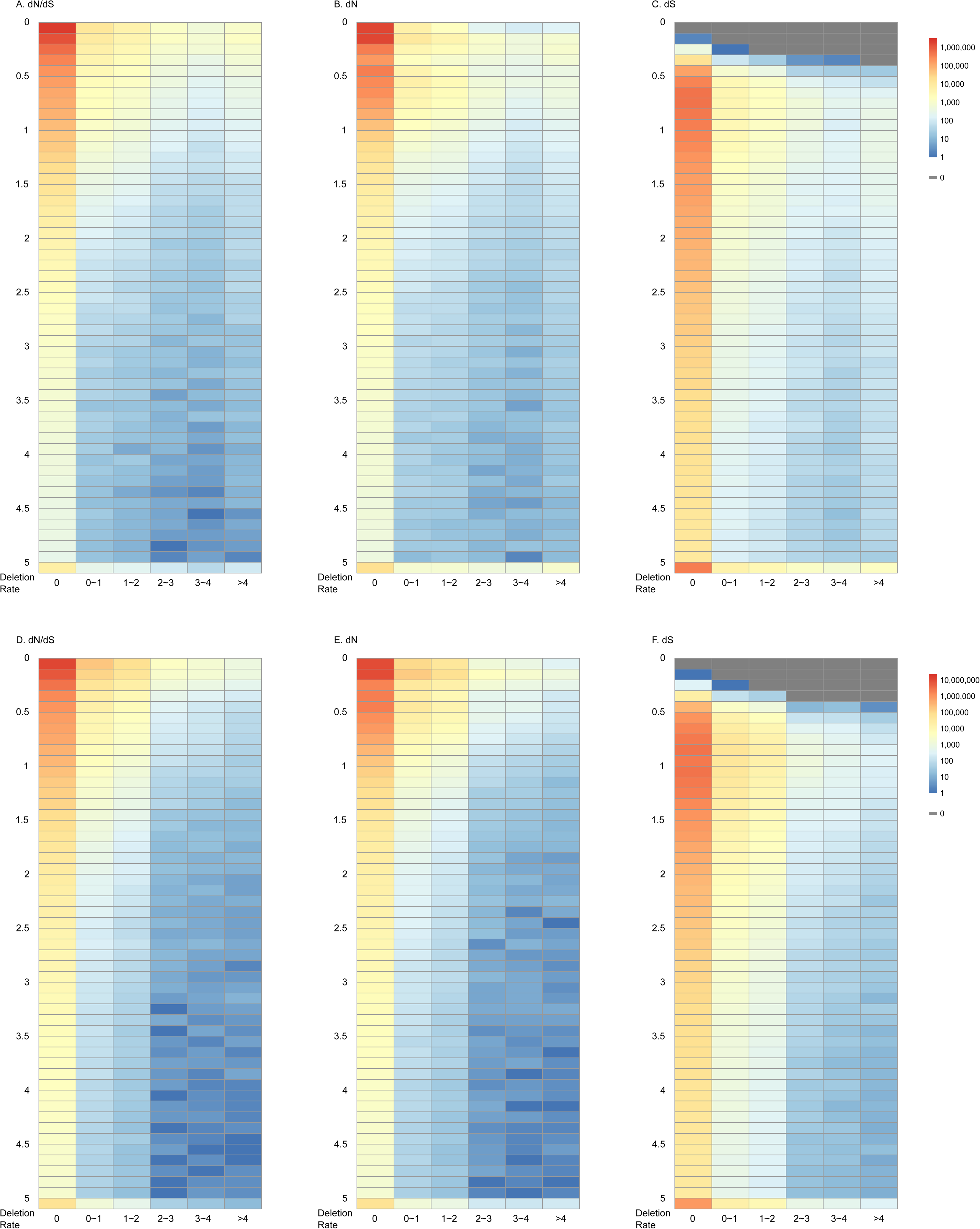
Density heatmap showing joint distribution of substitution measures and deletion rate in “All” dataset. **A.** Real data, dN/dS; **B.** Real data, dN; **C.** Real data, dS; **D.** Simulated data, dN/dS; **E.** Simulated data, dN; **F.** Simulated data, dS.

Considering that the interordinal relationship in Laurasiatheria is not entirely resolved (see Discussion), we re-calculated deletion rate using two alternative trees and calculated Spearman correlation coefficients with the same methods (Fig. S3). The difference from calculations based on the main tree is negligible.

### Deleted sites show higher rates of nonsynonymous substitution

If the rates of deletion and substitution are positively correlated then sites found to be deleted in at least one taxon would be expected to show a higher rate of substitution than sites that are present in all taxa. Fig. 5 summarizes the results of an analysis testing this prediction. We used Cohen’s *D*, a measure of effect size (Cohen 1988). Cohen’s *D* is the ratio of the difference between two distributions’ means and their pooled standard deviation. *D* < 0.2 is considered a small effect size, while *D* > 0.5 is a medium or large effect size. As predicted from the correlation analyses, both dN and dN/dS show medium to large effect sizes in both “All” and “NC-4+” datasets of the real data, whereas the simulated datasets have very small effect sizes. The differences for dS are also statistically significant (*Z* = 50.93 for “All” and *Z* = 10.94 for “NC-4+”, *P* < 0.0001 in both cases), but much smaller in magnitude. Further investigation showed that *some* but not all such effects seen in simulated data are due to alignment errors (Fig. S4). While both dN and dS has a positive effect size in TRUE alignment, they are cancelled out when the ratio, dN/dS is used; in such case TRUE alignment shows non-significant effect size in both “All” and “NC-4+.”

**Fig. 5.**
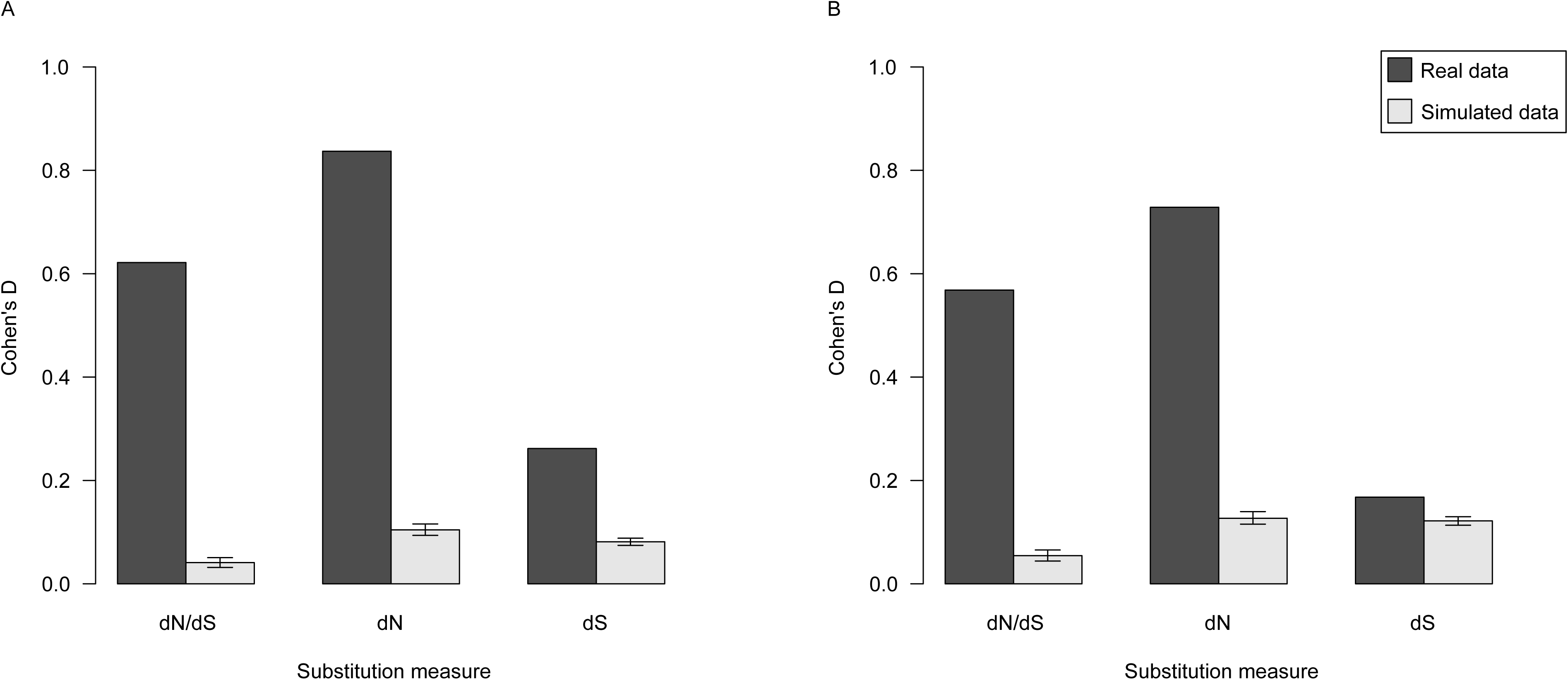
Effect size (Cohen’s D) indicating the difference of substitution measures (dN/dS, dN and dS) means between deleted and non-deleted sites. **A.** Based on the “All” dataset. **B.** Based on the “NC-4+” dataset, where all sites without any substitutions or present in less than four species were removed. For the simulated data, the shown value is the mean of 1,000 bootstrap re-samplings, and the error bars are 2.5% to 97.5% quantiles.

Fig. 6 compares the distributions of dN/dS in deleted and non-deleted sites in the “NC-4+” dataset. In the real data, 63.2% of deleted sites have dN/dS ≥ 0.2, while the number is 34.8% for non-deleted sites (Fig. 6A). The difference is negligible in the simulated data (Fig. 6B; 33.6% and 32.2% for deleted and non-deleted sites, respectively). A two-sample two-tailed Kolmogorov-Smirnov test (Kolmogorov 1933, Smirnov 1948) gives *D*_KS_ = 0.2886, *n* = 7.2×10^4^ & 3.6×10^6^ for real data and *D*_KS_ = 0.0262, *n* = 4.7×10^5^ & 2.2×10^7^ for simulated data (both *P* < 0.0001). These results confirm that rates of deletion and nonsynonymous substitution per site are positively correlated.

**Fig. 6.**
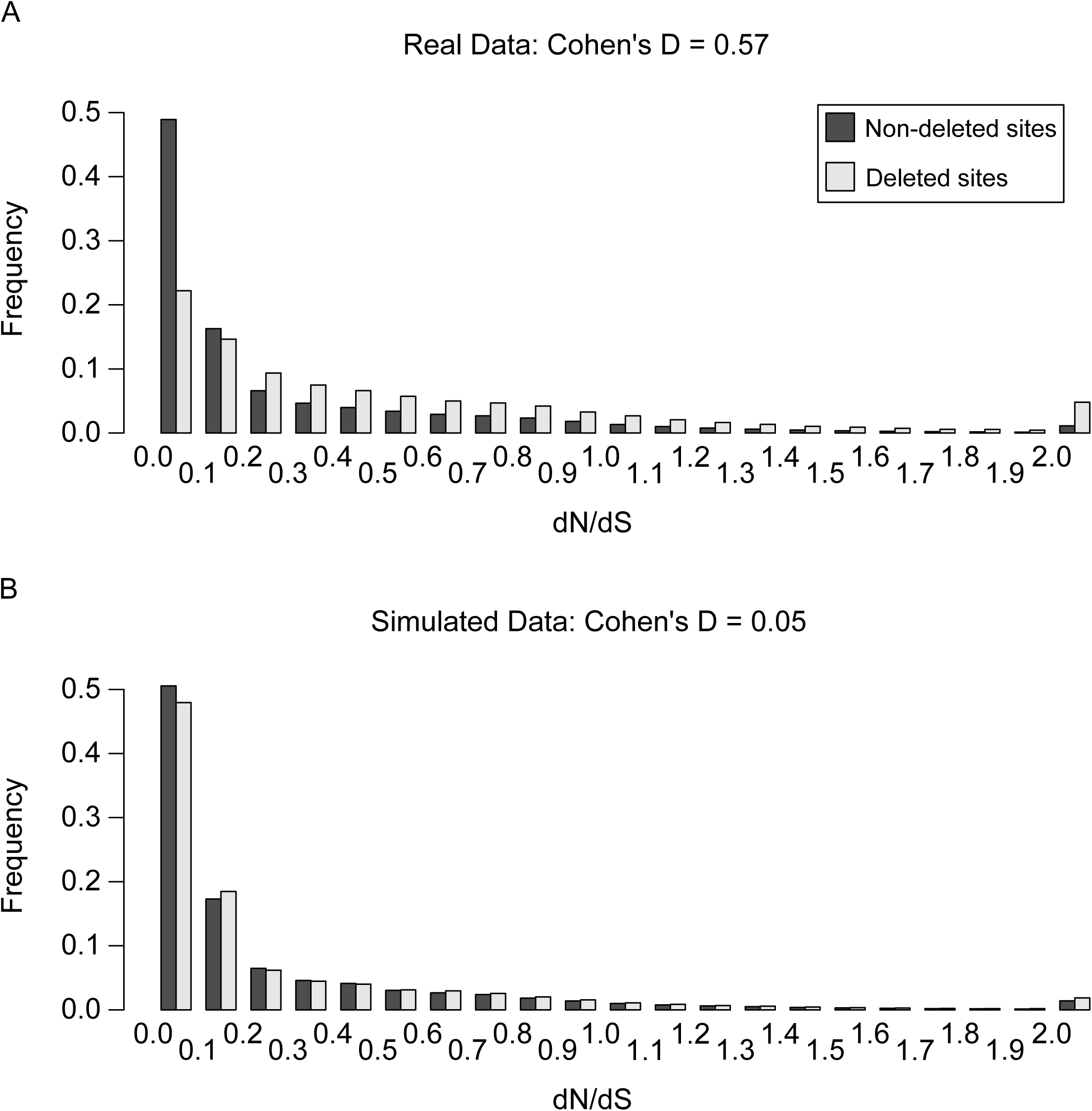
Histograms showing dN/dS distribution comparisons between sites with and without deletion, in both **A.** real and **B.** simulated data aligned with PROBCONS. The axis marks the lower bound of each bin, i.e. the bin marked “0” indicates 0≤dN/dS<0.1. It can be observed that the distributions are much more different in real data than in simulated data: the non-deleted sites have a heavier left tail, while the deleted sites have a heavier right tail.

### The deletion and nonsynonymous substitution rates per gene are positively correlated

To reduce the stochastic effects caused by limited number of mutations on each site, we decided to look at the same correlation at the gene level. We used the same statistical method with gene-averaged deletion rates and substitution measures.

As for the site data, the Spearman correlation coefficients between the deletion rate and substitution measures are significant and positive (Figs. 7 and 8; all *P* < 0.0001). However, the strength of correlation depends on the substitution measure used. For both dN and dN/dS, the correlation is strong (ρ ≈ 0.5), but for dS it is weak (ρ = 0.14). These correlations disappear completely in the simulated data (Fig. 8), and a negative but non-significant correlation is discovered with TRUE alignments of the simulated data (Fig. S5). Using a weighted deletion rate based on number of *codons deleted* and a rate based on number of *deletion events* does not seem to produce substantially different results (Fig. 8A and 8B), although the latter gives slightly higher correlation coefficients. We conclude that rates of deletion and nonsynonymous substitution per gene are positively and strongly correlated.

**Fig. 7.**
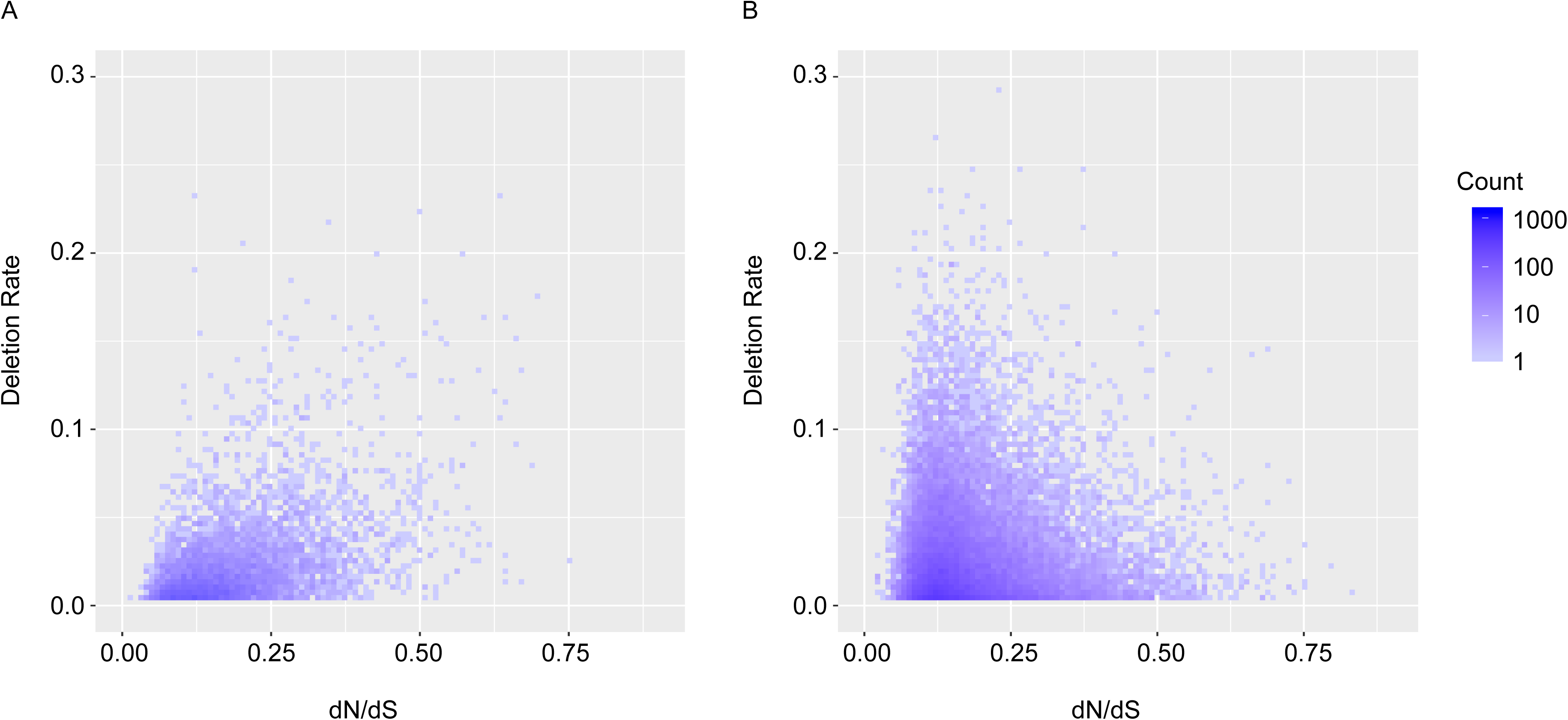
Gene-wise deletion rates plotted against dN/dS, in both **A.** real and **B.** simulated data. In real data, genes with high dN/dS (right) are more likely to have high deletion rate (up), which is not true in simulated data.

**Fig. 8.**
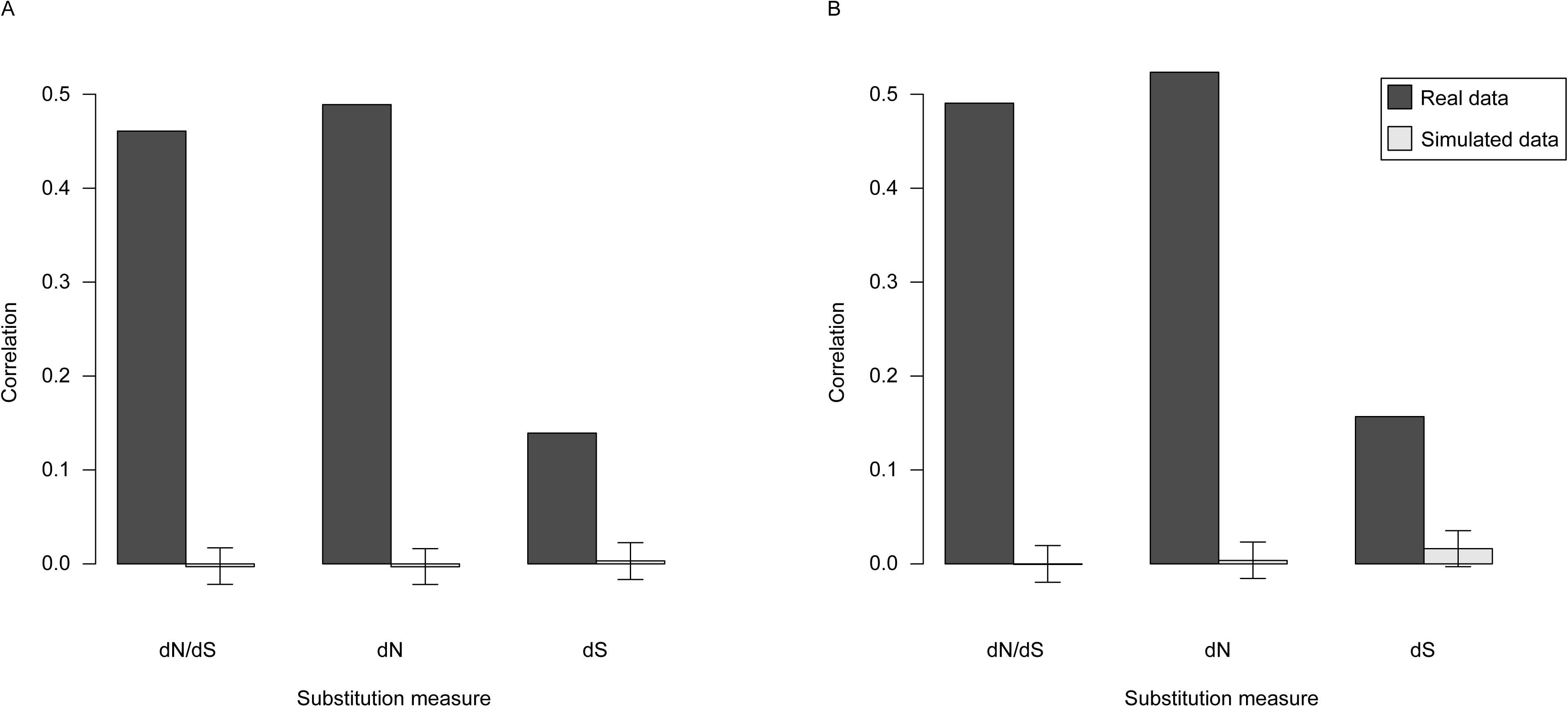
Gene-wise Spearman correlations between deletion rate and substitution (dN/dS, dN and dS) in real and simulated data. In both dN/dS and dN, the correlation in real data is very high (≈ 0.45) compared to simulated data (<0.05); the difference is much less pronounced in dS. For the simulated data, the shown value is the mean of 1,000 bootstrap re-samplings, and the error bars are 2.5% to 97.5% quantiles. The deletion rate was calculated based on **A.** number of codons deleted, **B.** number of deletion events.

### The deletion and nonsynonymous substitution rates within genes are positively correlated

We also analyzed the correlation within genes, to see whether the site-wise correlation is entirely caused by difference *between* genes. Fig. 9 shows the distribution of within-gene correlation for both real and simulated data. In real data, we only used 463 genes in “all” and 454 in “NC-4+” (Fig. S6) that have an estimated ancestral length over 1,500 aa and contains at least one deletion. In simulated data, 2,062 genes in “all” and 2,041 genes in “NC-4+” fit the same criteria and were used. In smaller genes the sample size is too small to generate reliable correlation coefficients. In dN/dS, the real data gives a slightly higher correlation compared to the simulated data (ρ ≈ 0.05 compared to ρ ≈ 0.02), although not to the level of genome-wide, site-wise correlation. dN produced a similar pattern.

**Fig. 9.**
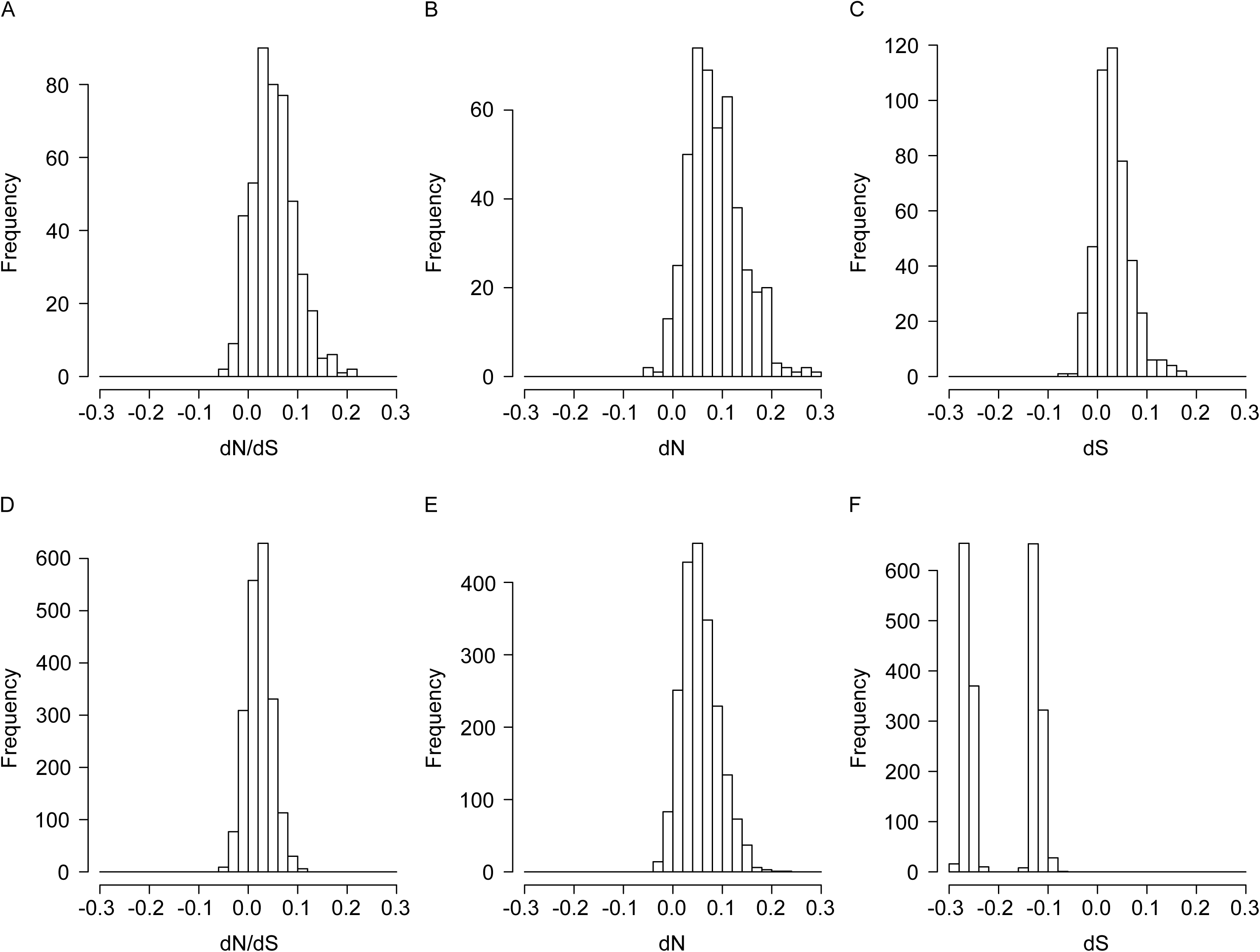
Histograms of distributions of within-gene Spearman correlation between substitution measures and deletion rate, using “All” dataset. Data are based on genes with an “ancestral” length of over 500 codons, and at least one deletion event. A total of 463 real genes and 2,062 simulated genes were used. **A.** Real data, dN/dS; **B.** Real data, dN; **C.** Real data, dS; **D.** Simulated data, dN/dS; **E.** Simulated data, dN; **F.** Simulated data, dS.

## Discussion

### Implications on protein sequence evolution

Our study shows that there is indeed a positive correlation between the probability of a codon being deleted and its dN/dS value (Figs. 3–8), indicating similarity in patterns of purifying selection against deletion and amino acid replacement. dN also produces a correlation to deletion rates, at a level similar to dN/dS. On the other hand, such correlation is very weak when dS is used, even undistinguishable from simulated data in some cases. This is unlikely because both types of mutation are correlated, because any mutation process affects dN and dS in the same way; instead, a more plausible explanation is that a common force, purifying selection, determines both replacement and deletion rates. This can be interpreted as meaning that both replacement and deletion can damage the function of an amino acid residue in the protein, thus reducing the fitness of individuals bearing such mutation. However, this site-wise correlation is weak, on the order of ρ ≈ 0.1; therefore, it would be difficult to predict one kind of selection from the other. In other words, selection against deletions is not completely consistent with selection against replacement.

We believe that one reason for the weakness of the correlation is the existence of “indifferent DNA” (Graur et al. 2013, 2015). Indifferent DNA refers to sequences that are subject to strong purifying selection against deletions but not substitutions, due to its functionality relies more on the length rather than the exact sequences. For example, it is possible that certain amino acids are required to maintain the spatial relationships between other amino acids in the protein and, therefore, cannot be deleted, but can be replaced by multiple amino acids with similar biochemical properties. Consistent with this idea, the scatter plot in Fig. 7 shows many genes with low deletion rate and high dN/dS, but few genes with high deletion rate and low dN/dS.

Our study on the correlation between substitutions and indels is the first one that involves genomic protein-coding genes, and includes both site-wise and gene-wise analyses. Using the deletion rate inferred from multiple sequence alignments instead of data on genetic diseases (Miller et al. 2007) made the rate estimation across multiple species rather than human-specific. Alignment-derived deletion rates are also available as long as the genomes of these species are annotated, while disease-derived rates are limited to clinical data and lethal sites are excluded. However, due to alignment errors and partial sequences in some species, alignment-derived deletion rates are less reliable. Nevertheless, we believe that we have taken precautions for these disadvantages, respectively by use of simulation and datasets “4+”/“6+.”

The potential non-independence between selection against substitutions and deletions can also be relevant in studies involving simulated sequence evolution. In protein simulation, the algorithm writer must decide whether to account for this correlation. For example, INDELible, one of the most comprehensive and frequently used simulation programs, does not allow variation of indel rates along the sequence (Fletcher and Yang 2009). On the other hand, programs like SIMPROT (Pang et al. 2005) implements an algorithm that chooses indel positions relative to their substitution rates. ROSE (Stoye et al. 1998) and indel-Seq-Gen (Strope et al. 2009) limit indels to less conserved regions of sequences.

### Difference between site-wise, gene-wise and within-gene analyses

Site-wise and gene-wise analyses on evolutionary parameters often yield different results (e.g., Wang et al. 2013). Here, we have shown that the Spearman correlation between dN, dS as well as dN/dS and deletion rate are much higher in gene-wise comparisons (Fig. 8) than in site-wise comparisons (Fig. 3). dN/dS values vary in a much larger range in site-wise than gene-wise analyses (Lindblad-Toh et al. 2011). The elevated non-synonymous substitution and deletion rates we observed are mostly due to relaxed purifying selection, but it is possible for a tiny minority of sites (individual amino acids) to undergo positive selection which yields a dN/dS above 1, causing a negligible effect on the correlation. This is rare for a whole gene because a protein’s basic structure need to be kept consistent for it to function, and it is almost impossible for the gene-wise signal to be caused by positive selection. Therefore, a site-wise study can provide a higher resolution on the selection schemes on the coding part of genomes. On the other hand, site-wise studies suffer from a low sample size for each data point, and thus larger sampling error and risk of being over-parameterized (Rodrigue et al. 2010).

The difference in the magnitudes of the gene-wise and site-wise correlations indicates that the gene-wise correlation is not entirely explained by site-wise correlations within genes. One possible mechanism for this discrepancy are differences in levels of selective constraint between proteins. Such differences would be expected to cause a positive correlation among genes that would not be detectable within genes.

An earlier study showed that most indels occur in intrinsically disordered regions of proteins, which are fast-evolving compared to structured regions (Light et al. 2013). A protein often contains both structured and disordered regions, thus this correlation would be present in within-gene comparisons, which is consistent with our results (Fig. 9).

### Artifactual correlation caused by alignment errors

Aside from the biological insights into protein sequence evolution, this study also provides information about consequences of alignment errors. There is no pre-determined correlation between indels and dN/dS in the simulated sequences, thus all estimated correlation is due to artifacts. The correlation between dN/dS and deletion in true alignments of simulated sequences is indistinguishable from zero, which confirmed this point. The same correlations estimated from inferred alignment, on the other hand, are consistently higher than zero. The only difference between them is the presence of alignment error, therefore we can conclude that the small correlation observed in simulated reconstructed alignments is caused by alignment errors.

Multiple sequence alignment is a mathematically difficult (NP-complete) problem. While an optimal solution exists theoretically, it cannot be computed within feasible time. All current multiple sequence alignment algorithms use heuristic methods. These algorithms typically produce alignments that are shorter than the true alignment due to preferring mismatches over gaps, and gives mathematically optimal placements while the real process is sub- or co-optimal (Landan and Graur 2008, 2009). Regions that are rich in insertions and deletions are difficult to align due to co-optimal placement of gaps, thus putting gaps and mismatches together more often than it should be.

On the other hand, there is a correlation between dN and deletion as well as dS and deletion in simulated sequences that cannot be explained by alignment errors. This phenomenon appears in both true and inferred alignments, and in both site-wise and gene-wise analyses. The most likely explanation is different rates of evolution (tree length) among different genes, because dN, dS and deletion rate are all indicators of total evolutionary change along the entire tree.

### Phase-1 and Phase-2 deletions

A phase-1 or phase-2 codon deletion (deletions that only partially encompass the first and the last codon involved) can cause an amino acid mismatch without nucleotide substitutions. They are also called non-conservative deletions because they do not conserve the undeleted amino acids (de la Chaux et al. 2007). However, past studies demonstrated that such events are less common than expected by chance. In a study on pairwise indel event between mouse and rat, 12% of indels found are non-conservative, in contrast with a simulation expectation of 29% (Taylor et al. 2004); another study (de la Chaux et al. 2007) gave an even lower estimate that 4% of all deletions are non-conservative from 3-primate alignments.

Unfortunately, with the simulation and alignment methods we used, we could not account for the effects for such deletions, nor could we mimic them by simulation. Nevertheless, the mismatch caused by non-conservative deletions usually does not happen in the same site as the gap. For example, if ACGCAT (Thr-His) became A---AT (Asn), the Asn residue will be aligned into one of the sites, while the gap occupies the other. The elevated dN/dS would thus only occur in the non-gap site. It is possible that the presence of such a mismatch complicates the alignment process and attracts other alignment errors, but we are not able to quantify this effect.

### Long deletions

Our study limited the length of deletion to 8 amino acids (24 nucleotides) or less. There are several reasons for excluding longer deletions. First, long indels in protein sequences usually accompany large changes in the protein’s function or structure. Repeatable protein structures such as alpha helix (Scholtz and Baldwin 1992) and zinc finger (Klug and Rhodes 1987) are usually ten amino acids or longer. Such large-scale changes in protein structure usually result in strong fitness effects and must be studied with a case-by-case basis and integrated with biochemical experiments. While short indels can have consequences in protein structural domains, they are usually preserved only in regions with weak purifying selection and do not change the protein’s function drastically (Zhang et al. 2011). Second, long gaps that can be interpreted as long deletions can co-occur with alignment difficulties. This includes, again, two situations: (1) Real long deletions can cause alignment errors because of unrealistic values of gap-extending penalties. (2) When highly diverged or non-homologous regions are aligned with each other, long gaps can occur as algorithmic artifacts. Non-homologous sections can exist in corresponding regions of orthologous proteins if structural mutations such as translocation occurred.

### Caveats and future directions

In our study, the simulation part was used as a negative control. In other words, it was used as a baseline when indel rates and dN/dS are independent from each other. We suggest that in future studies, a positive control can be implemented. If a simulation includes a correlation between indel and substitution models (or even perfectly linearly correlated rates), we could see how the results would compare to the real data. After all, even if the input indel and replacement rates are perfectly linear to each other, the site-wise correlation would still not be one because of stochastic effects.

In a neutral indel model by Lunter et al. (2006), the length of intergap segments (IGSs), gap-free regions of an alignment between two indel events, was identified as an important parameter. If indels were randomly distributed, IGS lengths would have a geometric distribution; instead, from a human-mouse comparison, this is only true for segments shorter than 50 bp. However, long IGSs (100 bp or more) are highly overrepresented than the expectation, indicating blocks that are resistant to indels, likely due to purifying selection. The model used in our study did not explicitly include the length of indel-free regions; in future studies it may be interesting to see which genes have the longest IGSs and how they correspond to substitution measures. Sampling of additional species would be useful to distinguish IGSs caused by purifying selection instead of stochastic effects.

In the phylogenetic tree used in this study, we put the horse (Perissodactyla) and the dog (Carnivora) together as sister groups, while the cow (Cetartiodactyla) is a sister group for the horse+dog clade. This hypothesis of Laurasiatherian evolution, known as Pegasoferae, is supported by a phylogenetic study using molecular data (Nishihara et al. 2006). However, the evolutionary relationship among horse, dog and cow is still under debate. A rival hypothesis groups the horse and the cow together (Perissodactyla + Cetartiodactyla = Euungulata), to the exclusion of the dog (Prasad et al. 2008). We have partially addressed the problem by re-calculating deletion rates using alternative trees and the change of results is negligible (Fig. S3). However, the FUBAR analysis and production of simulated data are all based on the Pegasoferae tree, and cannot be redone with other trees due to time constraints. We reasoned that in the rivaling hypotheses, the branch separating two of them from the third is very short, and this controversy would have a minimal effect on the estimation of evolutionary parameters. Therefore, we have arbitrarily chosen the Pegasoferae hypothesis. It may be a good idea to check if the choice of phylogenetic tree will affect the result in the future.

Incomplete lineage sorting (ILS) occurs when gene tree differs from the species tree (Maddison 1997), and introduces errors to any analyses based on phylogenetic trees. It is more likely to occur when two or more speciation events occur relatively close to each other. In our nine-species tree, the group that is most likely to suffer from such effect is Laurasiatherians (Hallström et al. 2011), but ILS occurring in other branches cannot be ruled out. We did not account for gene tree heterogeneity due to computational simplicity, but it may be a potential problem that could be resolved in future studies. Nevertheless, at least within Laurasiatheria, the use of alternative trees does not change our results in any meaningful way.

Finally, we used only protein-coding sequences in our study, because dN/dS, a reliable estimator of phylogenetic-level constraint, is only possible in protein-coding sequences. Selection against indels and substitutions in non-coding regions can be more efficiently studied in population-level analyses or between closely related species but this would be beyond the scope of this study. A future direction could be the extension of our conclusions into non-coding DNA sequences, especially in RNA genes.

### Conclusion

This study has demonstrated that in the evolution of mammalian proteins, the selection regimes on amino acid replacement and on short deletions are weakly correlated to each other. Codons that are less likely to undergo nonsynonymous substitutions are statistically also less likely to be deleted. However, in practice this correlation can be overestimated due to the effects of alignment errors.

## Materials and Methods

### Data collection and analysis of dN, dS and dN/dS

A list of aligned mammalian protein sequences was taken from Lindblad-Toh et al. (2011). To make sure that only good-quality genome sequences were used, we only included data from 9 mammalian species (Fig. 1): human (*Homo sapiens*), chimpanzee (*Pan tryglodytes*), macaque (*Macaca mulatta*), rat (*Rattus norvegicus*), mouse (*Mus musculus*), guinea pig (*Cavia porcellus*), dog (*Canis lupus familiaris*), cow (*Bos taurus*), and horse (*Equus caballus*). We retained 8,605 alignments.

Coding DNA sequences that correspond to these sequences were retrieved from ENSEMBL 2011 archive (Flicek et al. 2011).

All protein sequences were aligned with PROBCONS with default parameters (Do et al. 2005), and DNA sequences were aligned using the protein alignments as guides. Maximum likelihood trees were produced with RAxML (Stamatakis 2006) from the alignments, with a GTR+Gamma model and tree topology restricted to that of Fig. 1, and all other parameters set to default (standard hill-climbing algorithm). To reduce bias caused by unrealistic trees, 10 genes that produced a total tree length above 5 were discarded. (In the 8,605 genes, the mean tree length is 0.744 and standard deviation is 1.289. The shortest removed tree length is 6.897 and longest retained is 4.038.) Throughout the study, we used the remaining 8,595 genes. This correspond to ~42% of all human protein-coding genes. In phylogenomic studies, a trade-off between number of species and number of genes are well-noted; we decided on these nine species because of they represent all main branches of Boreoeutheria, which contains the vast majority of mammal species; these are also among the best annotated and highest quality genomes.

The DNA alignments were processed through the program HyPhy using the FUBAR script (Murrell et al. 2013), which estimated the dN and dS of each site using an approximate Bayesian algorithm, a Markov chain Monte Carlo process that compares a large number of site classes to identify and estimate selection. Their ratio ω = dN/dS was calculated from the output of FUBAR.

### Deletion identification and statistical analysis

Deletions of 1 to 8 amino acids were identified along seven pairs of branches (Fig. 1). These branch pairs are: (A) human and chimpanzee lineages (red branches, macaque as outgroup); (B) ape and macaque lineages (green branches, cow as outgroup); (C) rat and mouse lineages (indigo branches, guinea pig as outgroup); (D) murid and guinea pig lineages (orange branches, human as outgroup); (E) primates and rodents lineages (purple branches, cow as outgroup); (F) dog and horse lineages (yellow branches, cow as outgroup); (G) (dog+horse) and cow lineages (cyan branches, human as outgroup). The outgroup was used to determine whether a gap in the alignment is caused by an insertion or a deletion (Fig. 2A). In branch pair (B), the closest outgroup is a rodent, but cow was chosen because rodents have long branch lengths. For a lineage containing multiple species (e.g., apes), only the branch before the divergence (e.g., divergence between human and chimpanzee) was analyzed. This was done by combining multiple sequences into an “ancestral” sequence: any site that is a gap in *all* combined species is a gap site in the “ancestor”, and if the site is not a gap in at least one of these sequences, it is considered non-gap in the “ancestor.” In this way, every branch in the nine-species tree, excluding the root branch, was searched for deletions without repetition. The root branch (the branch separating the primates-rodents group and other mammals) was not searched for deletions because the directions of its indels could not be determined.

A fraction of amino acid sites are excluded from analysis because of ambiguity and difficulties in detecting deletions or substitutions. These sites include gaps in the outgroup (Fig. 2B), gaps in both ingroup taxa (Fig. 2C), deletions over 8 amino acids long (Fig. 2D), ambiguous amino acids (Fig. 2E), and terminal gaps (Fig. 2F). In some cases we excluded a site in the analysis of one lineage pair but not another.

The weighted deletion rate of an amino acid site, *D*, is calculated as *D* = 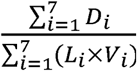 *D_i_* = 1 if that site is part of a deletion in the *i*th lineage pair, and 0 otherwise; *V*_*i*_ = 1 if that site is **not** excluded in that lineage pair, and 0 otherwise; *L*_*i*_ is the sum of branch lengths of the *i*th lineage pair, based on the placental tree (without chromosome X) from human/hg19/GRCh37 46 species multiple alignment (http://genomewiki.ucsc.edu/index.php/Human/hg19/GRCh37_46-way_multiple_alignment, Kent et al. 2002; Fig. 1).

Site-wise weighted deletion rates were re-calculated using two alternative trees that differ from the main tree in the relationship within Laurasiatheria; in one tree the horse and the cow were considered sister groups (Euungulata) and in the other the dog and the cow were considered sister groups. Because the branch lengths were not available for alternative trees, we used an *ad hoc* approach that kept the length of terminal branches and used the length of the internal branch (the one separating the horse-dog ancestor from the Laurasiatheria ancestor) for the new internal branches. This has minimal effects on deletion rate estimation because this branch is very short.

The weighted deletion rate of a gene, *D*_*G*_ is calculated as 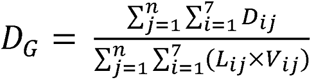 where *n* is the number of codons in the gene, *D*_*ij*_ is *D*_*i*_ in the *j*th codon in that gene, and *L*_*ij*_ is *L*_*i*_ in the *j*th codon in that gene.

An alternative gene-wise deletion rate is calculated as 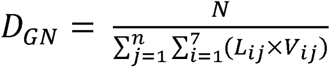 where *N* is the number of deletion events identified in any lineage in that gene. *D*_*GN*_ is called the event-number deletion rate of a gene.

For each amino acid site in each alignment, its deletion rate and three substitution measures (dN, dS and dN/dS) were obtained. For each alignment method, Spearman correlation coefficients were calculated between the weighted deletion rate, D, and the three substitution measures. This dataset uses all sites and is thus named “**All**.” See Table 1 for summary statistics on this dataset.

**Table 1.** A summary of our data, both real and simulated, based on “All” dataset. The Simulated Data was analyzed separately for the PROBCONS realignment and the true alignment. Cohen’s Ds were calculated for some statistics between real and simulated data aligned with PROBCONS to quantify the similarity between the two datasets.

|  | Real Data<br>(PROBCONS) | Simulated Data<br>(PROBCONS) | Cohen’s D between<br>real and simulated<br>data (PROBCONS) | Simulated Data<br>(TRUE) |
| --- | --- | --- | --- | --- |
| Number of genes | 8,595 | 42,975 | N/A | 42,975 |
| Total alignment length (aa) | 5675396 | 28458331 | N/A | 28465275 |
| Proportion of constant sites | 0.3106 | 0.2266 | N/A | 0.2275 |
| Number of deletions | 50698 | 338330 | N/A | 340895 |
| Mean deletion size (aa) | 1.9591 (1.5845) | 1.9218 (1.5434) | 0.0238 | 1.9274 (1.5412) |
| Mean site-wise dN (sd) | 0.3326 (0.9098) | 0.2804 (0.5671) | 0.0689 | 0.2793 (0.5625) |
| Mean site-wise dS (sd) | 1.9526 (2.6370) | 1.5625 (1.8169) | 0.1722 | 1.5586 (1.8032) |
| Mean site-wise dN/dS (sd) | 0.2687 (0.4891) | 0.2705 (0.5630) | 0.0034 | 0.2705 (0.5654) |
| Mean site-wise deletion<br>rate (sd) | 0.0353 (0.8410) | 0.0246 (0.2115) | 0.0174 | 0.0245 (0.1862) |
| Mean gene-wise dN (sd) | 0.3595 (0.2087) | 0.2983 (0.1432) | 0.3419 | 0.2974 (0.1433) |
| Mean gene-wise dS (sd) | 2.1206 (0.4309) | 1.6833 (0.2955) | 1.1836 | 1.6789 (0.2936) |
| Mean gene-wise dN/dS<br>(sd) | 0.1702 (0.0940) | 0.1790 (0.0884) | 0.0964 | 0.1789 (0.0888) |
| Mean gene-wise deletion<br>rate (sd) | 0.0166 (0.0222) | 0.0186 (0.0258) | 0.0831 | 0.0189 (0.0266) |

To reduce the effects of spuriously high or low values of dN/dS due to “gappy” sites, the correlation coefficients were recalculated for (1) sites that have not experienced a gap event in at least four sequences, and (2) sites that have not experienced a gap event in at least six sequences. These datasets are referred to as “**4+**” and “**6+**”, respectively. Many sites have not experienced any nucleotide substitution, and their dN/dS is technically incalculable due to division by 0, only approximated using extrapolation from other sites. Therefore, we generated sub-datasets in which these constant sites were excluded. These datasets were named “**NC-All**,” “**NC-4+**” and “**NC-6+**,” where “NC” stands for “no constant.”

### Coding sequence simulation and analysis

We simulated coding DNA sequences using INDELible (Fletcher and Yang 2009). INDELible evolves nucleotide sequences along the input tree based on a nucleotide substitution model. These substitutions are subject to selection as determined by dN/dS, randomly drawn from an input distribution for each site. Insertions and deletions, always multiples of three nucleotides, are independently modeled and have a uniform rate among sites; however, the number of indels is proportional to the branch length.

We simulated a total of 8,595 genes × 5 replicates. For each gene, the ancestral gene length and level of divergence were based on the values derived from the corresponding real gene (see Supplementary Text and Fig. S1 for details). The distribution of dN/dS was a gamma distribution with a shape parameter of α = 0.5 (approximated from real data) and a mean calculated from its real data counterpart. The distribution was discretized into 50 bins between 0 and 1 (0–0.02, 0.02– 0.04 …), 20 bins between 1 and 2 (1–1.05, 1.05–1.1 …) and 1 bin above 2. In each bin, the dN/dS value used was the median. If a bin (usually the ones with highest dN/dS) has a probability below 10^−6^ in the gamma distribution, it was not used. The absolute deletion rate for each gene was drawn from a gamma distribution with a shape parameter of α = 0.6 (approximated from real data) and mean = 0.79 (the mean S_AI_ from the real data), so that it is independent from substitution rate (see Supplementary Text); the relative indel rate was calculated based on absolute indel rate and branch lengths. Indel length was modeled with a power law distribution with the maximal length of 40 codons (Cartwright 2009).

The simulated protein sequences were aligned with PROBCONS (alternative alignment tools give identical results), and then nucleotide alignments were threaded through the protein alignments. We estimated deletion rates and substitution measures based on these alignments, as well as for the “true” alignment (as control for alignment error), as described above for real data. See Table 1 for summary statistics on the simulated data.

We used bootstrapping to generate plausible ranges of values of the sequence statistics to compare with the ones obtained from real data. We generated 1,000 bootstrap subsets of the simulated data. In each subset, one random replicate was chosen from the five for each of the 8,595 genes. Spearman correlation coefficients were calculated for each subset. Each subset was processed as described for real data to generate datasets of each type (“All,” “4+,” “6+,” “NC-All,” “NC-4+,” and “NC-6+”). For the Spearman correlation coefficients, the mean, standard deviation, and 2.5% and 97.5% quantiles were calculated. We used *Z*-tests to compare the Spearman coefficients derived from real and simulated data.

### Distribution of dN/dS in deleted sites

All real mammalian protein sites that have undergone at least one deletion in any lineage were extracted from the data set and their distributions of estimated dN/dS are computed. The distributions were compared with those from deletion-free sites with χ^2^ tests. Effect sizes (Cohen’s *D*, Cohen 1988) were calculated between dN/dS distributions in deletion and non-deletion codons. These analyses were only done on “All” and “NC-4+” datasets as representative of all six datasets. These procedures were repeated for the simulated data. Similar to the previous section, 1,000 bootstrap subsets were used, and the mean, standard deviation, and 2.5% and 97.5% quantiles were calculated. Z-tests were used to compare real to simulated data.

### Analysis of gene-wise and within-gene correlations

For both real and simulated data, we calculated gene-wise dN, dS, dN/dS and deletion rate. Gene-wise dN and dS are the mean of corresponding values of “4+” sites over the whole gene. We did not use “NC-4+” because excluding substitution-free sites is likely to lead to overestimation of the substitution measures. Gene-wise dN/dS is gene-wise dN divided by gene-wise dS. The calculation of two alternative gene-wise deletion rates, *D*_*G*_ and *D*_*GN*_, is described in a previous sub-section.

We calculated the Spearman correlation between gene-level deletion rate and substitution measures in both real and simulated data. Similar to previous sections, in the simulated data bootstrapping is used. Each subsample includes only one replicate for every simulated gene. The mean, standard deviation, and 2.5% and 97.5% quantiles were calculated. Z-tests were used to compare real to simulated data.

We calculated within-gene Spearman correlation between deletion rates and substitution measures, using 466 real genes and 466 × 5 = 2,330 simulated genes that have the derived “ancestral gene length” longer than 1,500 amino acids. The correlation coefficients are calculated for both “all” and “NC-4+” datasets. For the real data, genes (three such genes in “all” and twelve in “NC-4+”) that do not have any deletions identified were removed from the data, while the rest (463 in “all” and 454 in “NC-4+”) were used to calculate the mean and standard deviation.

## Acknowledgements

We used the Maxwell cluster from the Center of Advanced Computing and Data Systems (CACDS) at the University of Houston. CACDS staff provided technical support. We would like to thank Sarah Parks and her colleagues at EMBL-European Bioinformatics Institute for their help in running the SLR program on part of our data. R.B.R.A. was funded by NIH R01GM101352. We would also like to thank Jaanus Suurväli and Jan Gravemeyer at University of Cologne for their help in manuscript editing.

## Data availability

We uploaded our real and simulated alignments as well as Perl scripts of key steps on GitHub project “Mammal-Protein-Selection” (https://github.com/y-zheng/Mammal-Protein-Selection).

## Supplementary Files

Supplementary File 1. Contains Supplementary Text and Supplementary Figures 1-6.

Supplementary Text: Preliminary rounds of simulations to obtain simulation parameters Supplementary Figure 1: Flowchart describing the derivation and application of simulation parameters

Supplementary Figure 2: Comparison between reconstructed and true alignment: Spearman correlation between site-wise deletion rate and substitution measures

Supplementary Figure 3: Spearman correlation between deletion rate and substitution measures (dN/dS, dN and dS) in real and simulated data, with the deletions detected using alternative tree topologies regarding the internal relationship of Laurasiatheria.

Supplementary Figure 4: Comparison between reconstructed and true alignment: Cohen’s D between substitution measures in deleted and nondeleted sites

Supplementary Figure 5. Comparison between reconstructed and true alignment: Spearman correlation between gene-wise deletion rate and substitution measures

Supplementary Figure 6. Histograms of distributions of within-gene Spearman correlation between substitution measures and deletion rate, using “NC-4+” dataset.

## References

Ashburner M, Ball CA, Blake JA, Botstein D, Butler H, Cherry JM, Davis AP, Dolinski K, Dwight SS, Eppig JT, Harris MA. (2000) Gene Ontology: tool for the unification of biology. Nat Genet. 25(1):25.

Cartwright R. (2009) Problems and Solutions for Estimating Indel Rates and Length Distributions. Mol Biol Evol. 26:473–480.

Chen J-Q, Wu Y, Yang H, Bergelson J, Kreitman M, Tian D. (2009) Variation in the Ratio of Nucleotide Substitution and Indel Rates across Genomes in Mammals and Bacteria. Mol Biol Evol 26:1523–1531.

Cohen J. (1988) Statistical Power Analysis for the Behavioral Sciences (second ed.). Lawrence Erlbaum Associates. 67.

de la Chaux N, Messer PW, Arndt PF. (2007) DNA indels in coding regions reveal selective constraints on protein evolution in the human lineage. BMC Evol Biol 7:19.

Do CB, Mahabhashyam MSP, Brudno M, Batzoglou S. (2005) ProbCons: Probabilistic consistency-based multiple sequence alignment. ProbCons: Probabilistic consistency-based multiple sequence alignment. Genome Res 15:330–340.

Fletcher W, Yang Z. (2009) INDELible: A Flexible Simulator of Biological Sequence Evolution. Mol Biol Evol 26:1879–1888.

Flicek P, Amode MR, Barrell D, Beal K, Brent S, Chen Y, Clapham P, Coates C, Fairley S, Fitzgerald S, et al. (2011) Ensembl 2011. Nucleic Acids Res 39:D800–D806.

Graur D. (2016) Molecular and Genome Evolution. Sinauer Associates, Sunderland, MA.

Graur D, Zheng Y, Azevedo RBR. (2015) An evolutionary classification of genomic function. Genome Biol Evol 7:642–645.

Graur D, Zheng Y, Price N, Azevedo RBR, Zufall RA, Elhaik E. (2013) On the Immortality of Television Sets: “Function” in the Human Genome According to the Evolution-Free Gospel of ENCODE. Genome Biol Evol 5:578–590.

Hallström BM, Schneider A, Zoller S, Janke A. (2011) A genomic approach to examine the complex evolution of laurasiatherian mammals. PLoS One. 6(12):e28199.

Kent WJ, Sugnet CW, Furey TS, Roskin KM, Pringle TH, Zahler AM, Haussler D. (2002) The human genome browser at UCSC. Genome Res 12:996–1006.

Klug A, Rhodes D. (1987) Zinc fingers: a novel protein fold for nucleic acid recognition. Cold Spring Harb Symp Quant Biol 52:473–482.

Kolmogorov A (1933). Sulla determinazione empirica di una legge di distribuzione. G. Ist. Ital. Attuari. 4:83–91.

Landan G, Graur D. (2008) Local reliability measures from sets of co-optimal multiple sequence alignments. Pac Symp Biocomput 13:15–24.

Landan G, Graur D. (2009) Characterization of pairwise and multiple sequence alignment errors. Gene 441:141–147.

Light S, Sagit R, Ekman D, Elofsson A. (2013)Long indels are disordered: a study of disorder and indels in homologous eukaryotic proteins. Biochim. Biophys. Acta, Proteins Proteomics. 1834(5):890–897.

Lindblad-Toh K, Garber M, Zuk O, Lin MF, Parker BJ, Washietl S, Kheradpour P, Ernst J, Jordan G, Mauceli E, et al. (2011) A high-resolution map of human evolutionary constraint using 29 mammals. Nature 478: 476–482.

Lunter G, Ponting CP, Hein J. (2006) Genome-wide identification of human functional DNA using a neutral indel model. PLoS Comp Biol. 2(1):e5.

Lynch M. (2010) Evolution of the mutation rate. Trends Genet. 26(8):345–352.

Maddison WP. (1997) Gene trees in species trees. Syst Biol. 46(3):523–536.

Miller W, Rosenbloom K, Hardison RC, Hou M, Taylor J, Raney B, Burhans R, King DC, Baertsch R, Blankenberg D, et al. (2007) 28-Way vertebrate alignment and conservation track in the UCSC Genome Browser. Genome Res. 17:1797–1808.

Montgomery SB, Goode DL, Kvikstad E, Albers CA, Zhang ZD, Mu XJ, Ananda G, Howie B, Karczewski KJ, Smith KS, Anaya V. (2013) The origin, evolution, and functional impact of short insertion–deletion variants identified in 179 human genomes. Genome Res. 23(5):749–761.

Murrell B, Moola S, Mabona A, Weighill T, Sheward D, Pond SLK, Scheffler K. (2013) FUBAR: A Fast, Unconstrained Bayesian AppRoximation for Inferring Selection. Mol Biol Evol 30:1196–1205.

Nagy LG, Kocsubé S, Csanádi Z, Kovács GM, Petkovits T, Vágvölgyi C, Papp T. (2012) Re-mind the gap! Insertion–deletion data reveal neglected phylogenetic potential of the nuclear ribosomal internal transcribed spacer (ITS) of fungi. PloS One 7:e49794.

Nei M, Gojobori T. (1986) Simple methods for estimating the numbers of synonymous and nonsynonymous nucleotide substitutions. Mol Biol Evol 3:418–426.

Nishihara H, Hasegawa M, Okada N. (2006) Pegasoferae, an unexpected mammalian clade revealed by tracking ancient retroposon insertions. P Natl Acad Sci USA 103: 9929–9934.

Pang A, Smith AD, Nuin PAS, Tillier ERM. (2005) SIMPROT: Using an empirically determined indel distribution in simulations of protein evolution. BMC Bioinformatics 6:236.

Prasad AB, Allard MW, NISC Comparative Sequencing Program, Green EE. (2008) Confirming the phylogeny of mammals by use of large comparative sequence data sets. Mol Biol Evol 25:1795–1808.

Price N, Graur D. (2016) Are Synonymous Sites in Primates and Rodents Functionally Constrained? J Mol Evol 82:51–64.

Rodrigue N, Philippe H, Lartillot N. (2010) Mutation-selection models of coding sequence evolution with site-heterogeneous amino acid fitness profiles. Proc Natl Acad Sci USA 107(10):4629–4634.

Scholtz JM, Baldwin RL. (1992) The mechanism of alpha-helix formation by peptides. Annu Rev Biophys Biomol Struct 21(1):95–118.

Smirnov N (1948). Table for estimating the goodness of fit of empirical distributions. Ann Math Stat 19:279–281.

Stamatakis A. (2006) RAxML-VI-HPC: maximum likelihood-based phylogenetic analyses with thousands of taxa and mixed models. Bioinformatics 22:2688–2690.

Stoye J, Ever D, Meyer F. (1998) Rose: generating sequence families. Bioinformatics 14:157–163.

Strope CL, Abel K, Scott SD, Moriyama EN. (2009) Biological Sequence Simulation for Testing Complex Evolutionary Hypotheses: indel-Seq-Gen Version 2.0. Mol Biol Evol 26:2581–2593.

Sung W, Ackerman MS, Dillon MM, Platt TG, Fuqua C, Cooper VS, Lynch M. (2016) Evolution of the insertion-deletion mutation rate across the tree of life. G3: Genes, Genomes, Genetics. 6(8):2583–2591.

Sung W, Ackerman MS, Miller SF, Doak TG, Lynch M. (2012) Drift-barrier hypothesis and mutation-rate evolution. Proc Natl Acad Sci USA 109(45):18488–18492.

Taylor MS, Ponting CP, Copley RR. (2004) Occurrence and Consequences of Coding Sequence Insertions and Deletions in Mammalian Genomes. Genome Res 14:555–566.

Wang H, Susko E, Roger AJ. (2013) The Site-Wise Log-Likelihood Score is a Good Predictor of Genes under Positive Selection. J Mol Evol 76:280–294.

Waterston RH, Lindblad-Toh K, Birney E, Rogers J, Abril JF, Agarwal P, Agarwala R, Ainscough R, Alexandersson M, An P, et al. (2002) Initial sequencing and comparative analysis of the mouse genome. Nature 420:520–562.

Zhang Z, Huang J, Wang Z, Wang L, Gao P. (2011) Impact of indels on the flanking regions in structural domains. Mol Biol Evol 28(1):291–301.

